# NeuroTest: a benchtop testbed for evaluating sensing-capable electrophysiology and neurostimulation systems

**DOI:** 10.1101/2025.09.19.677167

**Authors:** Jeremiah K Morrow, Hanbin Cho, Moaad Benjaber, Timothy K Denison, Alik S Widge, Jeffrey A Herron

## Abstract

**Objective:** The development of increasingly complex implantable neuromodulation devices requires extensive benchtop validation and testing prior to *in vivo* implementation. Our goal is to enable this *in vitro* testing by creating a benchtop system that is accessible, easy-to-use, and functions reliably across academic and industry environments, reducing discordant results that arise from discordant approaches. Furthermore, such a system paves the way for standardizing benchtop characterization of future implantable systems by establishing a repeatable and comparable methodology for validating these devices and their capabilities.

**Approach:** We describe the NeuroTest Board (NTB), a low-cost, open-source device that combines a low-noise current source waveform generator with a high-precision data acquisition system. We provide information on its architecture, operation, and performance as an initial benchmark of the potential capabilities of the system.

**Main results:** We demonstrate the use of the NTB to assess electrophysiology systems in two separate applications. In one case, we assess a commonly used laboratory system (the RHD2132 headstage and Open Ephys data acquisition system), while the second case characterizes an investigational neuromodulation device in development for human use (the CorTec Brain InterChange). Following this initial sensing characterization, we explore stimulation-related artifacts that would hamper *in vivo* data collection. With the NTB, we develop and evaluate artifact mitigation strategies for each system, thus demonstrating an example application of this benchtop testbed.

**Significance:** The ability to test neuromodulation devices against a common benchmark will enable faster development of novel therapeutics by increasing inter-institutional reliability and decreasing troubleshooting time.

## 1. Introduction

As the field of neuromodulation research grows, there is a complementary rise in increasingly complex devices that expand the features and research capabilities available to scientists. Electrode arrays used for sensing and characterizing complicated neural dynamics continue to increase in density [1–3] and diversify in terms of materials and form factors [2,4,5], while adaptive stimulation paradigms are opening new therapeutic avenues for a range of neuropsychiatric disorders [6–12]. Adoption of new systems for use *in vivo* first requires thorough characterization and assessment of their functionalities [13]. However, verification of these systems is limited by the need to test with realistic electrophysiological signals in a controlled and consistent environment that mimics the tissue-electrode interface.

Initial feasibility and safety assessments of novel systems and approaches are often performed using preclinical models. While animal models enable biocompatibility assessments and scientific investigation *in vivo*, they also require significant investment (e.g., per diem costs, housing, investigator care/husbandry, etc.). Device development via animal model testing also often requires species-specific customization, for example, downscaling the size of a device, power supply, or connectors relative to the ultimate clinical design [14,15]. Those changes may limit the ability of animal models to predict the performance of a system when implanted in humans. Furthermore, *in vivo* conditions can introduce factors that may complicate system troubleshooting. The biggest issue being that it is difficult, or even impossible in some cases, to distinguish which aspects of incoming data are driven from a specific source of interest (e.g., neural activity in a target brain area) versus those that are a function of environmental noise, non-neural activity, or activity from a neural source the researcher is not interested in. This is particularly relevant when assessing how electrical stimulation alters activity in excitable tissue. Because stimulation is designed to elicit a reaction from the tissue, every pulse is thus accompanied by a mixture of stimulation-related artifact and potentially pertinent physiological responses. These artifacts affect not only real-time processing pipelines for closed-loop algorithms, but also the detection of relevant biomarkers such as stimulation evoked potentials (EPs). Specifically, determining what *in vivo* attributes (e.g., cellular responses or morphology) as opposed to what system characteristics may be responsible for unexpected outcomes can be a challenging confound. How to disentangle these components is an active and robust area of study [16–19].

The tools currently available to evaluate novel systems and adaptive algorithms are limited. Traditional laboratory equipment is constrained in its ability to simulate biopotential activity and many neuroscience labs lack the expertise to assemble the appropriate equipment in-house. Many function generators strictly generate standard electrical waveforms (e.g., sine or square waves). Arbitrary waveform generators can struggle to reproduce low-amplitude signals (i.e., below 1mV) typical of neural activity without the need for external attenuators, which are often custom made. Waveform generators that inherently generate low-amplitude signals tend to be more expensive, possibly beyond the reach of a laboratory’s budget for equipment that solely enables testing. Some proprietary signal generators [20,21] that allow for playback of simulated neuroelectric activity have facilitated benchtop evaluation with neural test signals. However, they present a limited set of signals available for playback (e.g., may come with only one preloaded neural recording), which may not model the electrophysiological characteristics of the intended *in vivo* application. These systems are also typically purpose-built around interfacing with a limited range of other devices, typically ones made by the same manufacturer. Lastly, standard commercial products for signal generation typically are not adaptive. The signal output can be manually updated by the user, but the devices themselves do not react to changing features of incoming data. This severely limits the ability of researchers to test novel technologies that rely on real time updating of stimulation parameters according to ongoing biological states. To expand on the features provided by these commercial solutions, more customizable simulators [22,23] and hardware-in-the-loop testing frameworks [24] have been introduced that provide multichannel, low-noise signal playback. These benchtop tools enable customized signal generation, greatly expanding the types of test signals to better capture the specific, target application of interest.

While these research tools expand the capabilities of benchtop evaluation, they also have limitations that should be considered. Most of the cited testbench solutions evaluate the sensing capabilities of neuromodulation devices either through direct voltage injection or a resistive load setup. These test systems fail to account for the effects of the tissue-electrode interface, particularly the impact of stimulation on this interface and the sensing circuit. This is a critical oversight. The nervous system functions as an electrolyte medium in which ions serve as the charge carriers, as opposed to the electrons that flow in resistive loads. The electrode-electrolyte interface adds an inherent capacitance and the potential for redox chemical reactions if voltage windows are exceeded. Both phenomena can filter and distort both applied stimulation waveforms and recorded tissue responses. Testing in an electrolyte, such as a saline bath, better mimics *in vivo* conditions. Lastly, testing within saline setups is considered standard procedure under the International Organization of Standards [25]. Accounting for these realities is critical when determining the performance of adaptive and closed-loop neural interfaces, especially when the ultimate goal is clinical use.

Another gap in existing benchtop testing environments is the lack of a means to assess whether a system can accurately measure the biological response to stimulation. The goal of neuromodulation systems is to change neural activity, and their sensing capabilities are often designed to provide closed-loop measurement to guide therapy. In most cases, however, we do not know how quickly the sensing components can detect a biological change, nor how large of a change is needed to enable reliable detection. As a non-limiting example, consider evoked potentials (EPs), which are a common metric for assessing connectivity [26–28] and neural excitability [29,30]. Their detection might be affected by whether the stimulated tissue can respond to the provided stimulation, e.g. by refractory periods. Aspects of the electrophysiology system, such as sensing configurations and stimulation artifact, may also influence the detection of an EP (e.g., long amplifier settling times obscure neural responses to stimulation). Additionally, signal processing techniques have been demonstrated to affect cortico-cortical evoked potentials (CCEPs) [31], further obfuscating what constitutes a “real” response. These factors present challenging confounds when attempting to debug a system and identify system characteristics that may be responsible for unexpected outcomes.

The above considerations establish needs for a system to evaluate and validate novel sensing/stimulation devices. The ideal test system would be designed around testing in saline (or other electrolyte) mediums using electrodes representative of those to be used in the implanted context (i.e., same size, materials, and form factor). Further, to properly test a bidirectional neural interface, the test system should be bidirectional, i.e. it should both sense device output and produce simulated neural data. Further, that simulated data should change in response to device output. Simulating neural signal changes due to stimulation adjustments is critical for evaluating the performance of proposed adaptive algorithms. Assessing the impact of stimulation artifact on the measurement of stimulation evoked activity simply cannot be achieved using a function or waveform generator. Instead, the test infrastructure for neural devices requires a bidirectional approach where the test system can be configured to adjust saline bath input based upon a model of how the nervous system would respond to the actions of the device under test. Lastly, the ideal system should be accessible in terms of price and ease-of-use and produce highly consistent results across test sites.

Here, we detail the NeuroTest platform, an *in vitro* benchtop testing system that supplements biopotential signal playback with its own recording capability and built-in waveform generation. A novel evoked potential mode enables stimulation evoked potential simulation, allowing researchers to conduct tests with reactive and/or closed-loop paradigms. These features allow for a more comprehensive benchtop characterization of the performance of novel neuromodulation technologies, helping users to be better informed about the capabilities of their target devices prior to approaching *in vivo* experiments. The system accelerates the process of designing and validating bidirectional implantable interfaces and can be used to create inter-laboratory standards for assessing system functionality. We describe the architecture, basic functions, and some simple applications of this platform for testing equipment used in neuromodulation research.

## 2. NeuroTest Board (NTB) Overview

The NeuroTest platform is a custom test bench setup designed to evaluate electrophysiology systems *in vitro* (Figure 1). It enables the characterization and testing of a device’s ability to sense signals in the presence of electrical stimulation in a saline-based environment. Evaluation within an *in vitro* environment enables controlled testing of the impact of the tissue-electrode interface on the device’s sensing circuit and assessment of how that impact may vary with stimulation.

**Figure 1:**
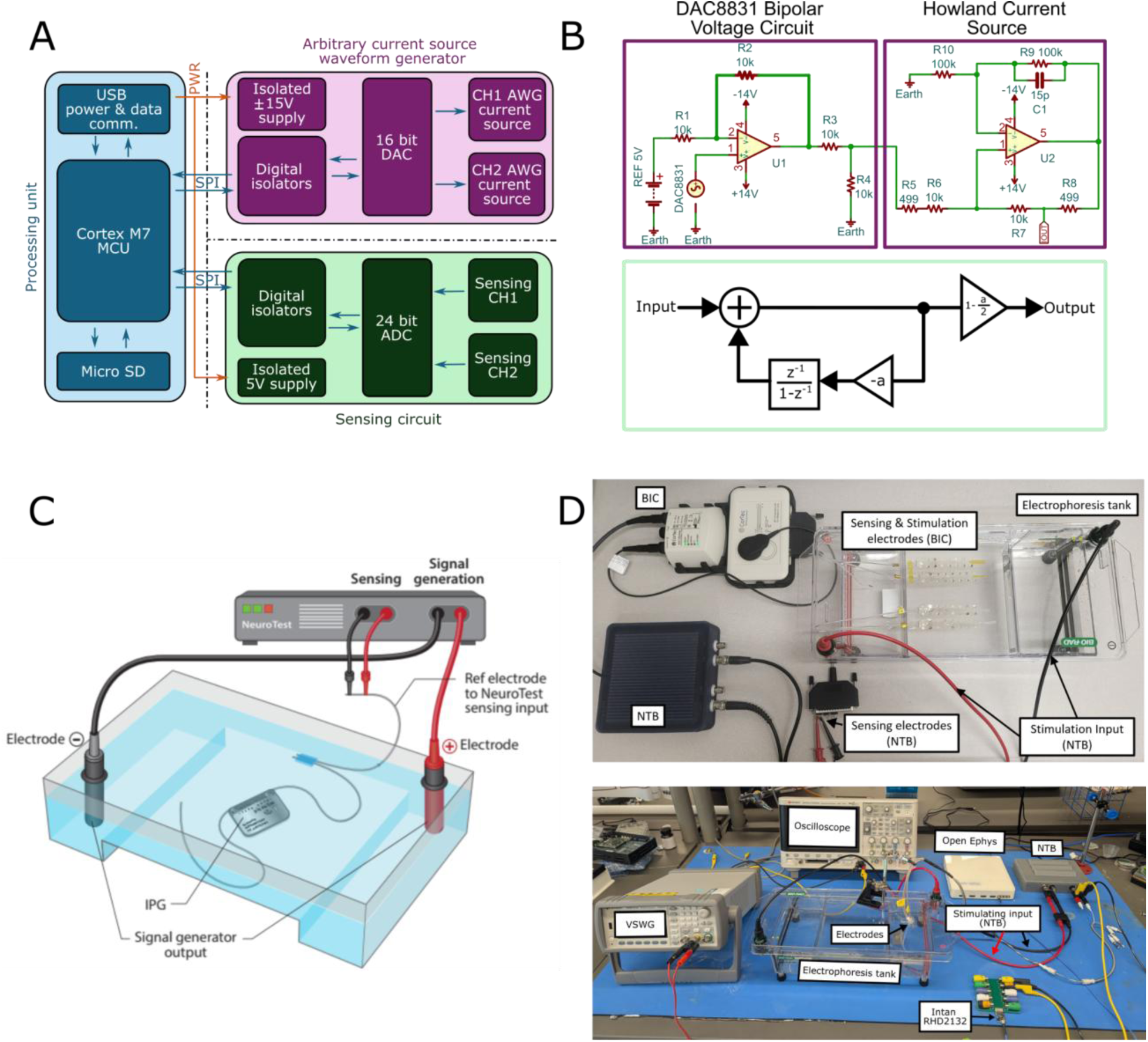
General setup overview. **(A)** Block diagram depicting the components of the NeuroTest Board (NTB). The system is divided into three components: the processing unit (blue), the waveform generator (purple), and the sensing circuit (green). Power lines (PWR) are shown in orange while communication between the components is shown by the blue arrows. Serial peripheral interface cables (SPI) are used for bi-directional communication between the processing unit and the sensing circuit and waveform generator. **(B)** Design for system components of the NTB. Color coding follows the format shown in A. (Top) Circuit schematics of the current source waveform generator to drive constant current over a range of load resistances. (Bottom) Programmable high pass filter topology integrated with the sensing circuit. **(C)** A cartoon depiction of the NTB (top) connected to a gel electrophoresis tank and a submerged implantable pulse generator (IPG) (bottom) **(D)** (Top) Picture of the benchtop setup at the clinical site. Sensing and stimulating electrodes from the CorTec Brain InterChange (BIC) device are submerged in 0.9% saline in the electrophoresis tank. The sensing side of the NTB is connected to two electrodes from an identical sEEG array using a modified D-sub connector. The waveform generator of the NTB is connected to the cathode and anode of the tank. (Bottom) Picture of the benchtop setup at the preclinical site. A Keysight 33500B voltage source waveform generator (VSWG) and a Keysight oscilloscope (DSOX2004A) were used to verify proper NTB function. Up to six electrodes (Pt-Ir, 125um diameter, 1-20kΩ) submerged in 0.9% saline are used for sensing or stimulation. Sensing electrodes are routed to the NTB (directly) or an Intan RHD2132 headstage (via a breadboard) using banana plugs. The signal from the same electrodes can be split and routed to both systems (as shown here, yellow and black cables in the NTB). An electrical stimulator (e.g., StimJim, not shown) can be connected to the electrodes, as well. The waveform generator of the NTB is connected across the cathode and anode of the tank similar to the clinical setup.

The NTB uses a stripped-down gel electrophoresis chamber (BioRad Inc) filled with a 0.9% saline solution in which the system under test is submerged. While an electrophoresis tank may not be strictly necessary, maintaining a stable and even electrical field across the test medium is crucial for repeatability. These tanks provide an accessible commercial product that many labs will be familiar with, making them a reasonable option for the testing medium. The tanks feature banana-plug connectors that are attached to wires submerged in the saline, which function as a cathode and an anode. Connecting these to a signal generator allows for voltage or current to be injected across the entire tank.

### 2.1 NTB Design

The electronics of the NTB were designed to generate simulated biopotential signals and to sense the output of a neural device under test. The NTB includes three main components: 1) a processing unit, 2) a precision arbitrary current source waveform generator (ACSWG), and 3) a precision sensing circuit. These three components of the NTB are galvanically isolated from each other to reduce noise, interferences, and ground loops between the different circuit functions. A system diagram of the NTB is illustrated in Figure 1A and additional technical schematics of its components are shown in Figure 1B.

#### 2.1.1 Processing unit

The first section of the NTB contains the microprocessor, a 32-bit Arm Cortex M7 clocked at 600 MHz, which is programmed to be controlled and powered with a USB 2.0 connection that enables communication speeds up to 480Mbps. This section of the board also includes a microSD card that allows users to load neural data for replay. This microSD card can also be used to save data observed during tests with the NTB, though data can also be saved directly to a control computer via the USB interface. The NTB is powered by a 5V supply via the USB connection. This connection also allows the user to update firmware on the NTB via the open-source Arduino IDE. Likewise, data can be streamed to the connected computer and visualized in the Serial Plot tool from Arduino or other data visualization software. The precision ACSWG and the precision sensing circuits are both powered from the same source as the processing unit, using two galvanically isolated DC-DC converters. One ±15V supply powers the ACSWG circuit, while the sensing circuit is supplied by an isolated 5V supply. All communication between the processor unit and the other two components is established using digital isolators.

#### 2.1.2 Precision arbitrary current source waveform generator

The NTB has two independent low-noise ACSWGs based on a Howland current source architecture [32] (Figure 1B). The current source circuit is designed to generate currents in the range of 0-10mA with a 305nA resolution driven by an ultra-low noise 16-bit digital-to-analog converter (DAC) combined with a low-noise, ultra-low offset amplifier. Although all sensing applications of a device under test will be dependent on electrode impedance and position of the recording dipole, the range and precision of the ACSWG allows a user to mimic a myriad of neuro-electric phenomenon in the scales observed *in vivo* [33]. For instance, LFP signals observed with current-generation sensing-capable implants tend to be in the 1-200 µV range [34,35], and *in vivo* stimulation evoked potentials in the 100-1000 µV range [36,37]. The ACSWG has a voltage range of ±14V supplied from low-noise linear regulators (LDOs) to enable the system to maintain the required current delivery over a large range of impedances depending on the *in vitro* setup used.

The bandwidth of the current source is reduced from 1MHz to 100kHz using a feedback capacitor in order to reduce the noise introduced by the NTB when connected in saline. Specifically, inadequate anti-aliasing by a device being tested can cause high frequency noise to fold back from the current source, distorting the sensing capabilities of the test device. Furthermore, we programmatically set the generator to have a range of 0.1 to 1000Hz by default to enable simulation of a wide range of neural oscillatory events [33], though this range can be changed by an end user as needed.

The output of the ACSWG can be tuned to produce voltages that are within expected ranges for the specific recording setup of an end user. That is, the amount of current being output is scaled such that the voltage values detected by a downstream data acquisition system (the “realized voltage”) are in the appropriate ranges when considering electrode size and material, dipole location, and other elements of the recording arrangement. This calibration is achieved via a function made available in the user interface. The function works by injecting sine waves into the saline tank at progressively increasing amplitudes via the ACSWG, which are then measured by the sensing component of the NTB. To ensure proper calibration, the system automatically adjusts the gain of the output such that the measured values more closely match the expected values.

#### 2.1.3 Precision signal acquisition

The NTB features a two-channel, high-precision data acquisition circuit with a 24-bit resolution delta-sigma analog-to-digital converter (ADC) supplied by an isolated 5V DC-DC converter. The data acquisition circuit has a theoretical noise floor of 1μV RMS at a 1kHz bandwidth, when gain is set to 32V/V and sampling frequency set to 2kHz. The data acquisition circuit also offers a configurable sampling frequency up to 32kHz, and configurable gain from 1 to 128V/V.

The precision sensing circuit has a programmable high pass filter built into the ADC that can be configured to filter out DC offsets and low frequency noise. The range of the high pass filter is dependent on the sampling frequency set of the ADC for the NTB. For example, with a 2kHz sampling rate, the high pass filter can be configured between 4mHz - 90Hz. The precision sensing circuit is controlled via the processing unit through digital isolators such that the sensing circuit is fully isolated from the other two parts of the NTB, reducing noise and crosstalk. The ADC sampling frequency of the NTB can be programmatically set up to 32kHz. However, as most bioelectrical field phenomena of interest occur between 0.1 and 500 Hz, the default value is set to 1kHz.

## 3. Benchtop Validation of NTB Functionality

Benchtop assessments were performed to evaluate the functionalities of the NTB. A diagram of an intended use setup with the NTB is visualized in Figure 1C. Considering the various settings used within the neuromodulation space, we designed the NTB to adapt to a wide variety of applications. To demonstrate this versatility and reliability, two distinct benchtop environments were configured for assessments at two separate locations. Testing at one site focused on preclinical applications (hereafter referred to as the “preclinical” site/setup) while evaluation done at the other site centered on clinical applications (hereafter referred to as the “clinical” site/setup). The preclinical benchtop setup sought to assess the characteristics of an Intan RHD2132 amplifier headstage (Intan Technologies) paired with the OpenEphys data acquisition system (Figure 1D). The clinical testing environment used the investigational CorTec Brain Interchange (BIC) system (Figure 1D).

### 3.1 Devices under Test

#### 3.1.1 Preclinical Benchtop Environment

In the preclinical setup (Figure 1D, bottom), between six and twelve platinum-iridium electrodes (built in-house using 125μm diameter wire from AM Systems, 1-20kΩ impedances) were submerged in the 0.9% saline bath. These electrodes were wired to banana plugs, which were used to connect to various downstream devices. Simple banana-to-BNC adapters were used to connect electrodes to the sensing side of the NTB, a StimJim electrical stimulator [38], or a Keysight digital oscilloscope (DSOX2004A, Keysight Technologies). A splitter was used to route parallel signals from the NTB sensing channel to a custom banana plug breakout board. This board was used to connect banana plugs to a 32-channel Omnetics connector, which in turn allowed the signal to be routed through the Intan RHD2132 headstage. The headstage was then connected to an Open Ephys DAQ [39] via a serial peripheral interface (SPI) cable. Either a Keysight voltage source waveform generator (33500B, Keysight Technologies) or the precision arbitrary current source waveform generator of the NTB was connected to the cathode and anode of the electrophoresis tank using banana-to-banana plug connectors. Both the oscilloscope and voltage source waveform generator provided parallel means to cross check that the various components of the preclinical setup were functioning as expected (e.g., we checked that the NTB, Open Ephys, and oscilloscope all registered sine waves injected via the NTB or waveform generator at the expected amplitudes and frequencies).

The Intan RHD2132 used in these tests features a 16-bit ADC and 32 unipolar amplifiers with integrated analog filters. We shorted the reference and ground pins together and verified that all headstages used were fully functional (i.e., no dead channels or unusual noise characteristics).

#### 3.1.2 Clinical Benchtop Environment

The benchtop environment with the BIC system consisted of two sets of 32 channel electrocorticography (ECoG) grids submerged in a saline solution. A single set of ECoG electrodes consisted of two grids of 2×8 platinum-iridium circular contacts and ended in a D-Sub connector. A set of ECoG grids were connected to the BIC and NTB each. While the D-Sub connector directly connected to the implant module of the BIC development kit, additional connectors were used to connect the NTB to specific pins of the D-Sub connector. This connection scheme made it such that only one channel was used for sensing with the NTB. The output from the NTB was connected to the cathode and anode of the saline tank. An additional ground electrode, an optional lead hard-wired to the implant casing connected to the ECoG grids, was also submerged in saline. However, the ground electrode was not shorted to any contacts on the ECoG grid. These channels were not shorted together during subsequent testing to more realistically reflect noise characteristics to be found *in vivo* conditions.

### 3.2 Process for Characterizing Noise and Artifact

To characterize system noise of the NTB, electrodes connected to the NTB and either the Intan RHD2132 or BIC were submerged in a saline solution. For these noise tests, a single sensing channel on the NTB was connected across a single pair of recording and reference electrodes that were submerged in the saline. Prior to recording, the NTB underwent calibration to ensure measurement reliability. Once calibration was completed, the NTB and both systems under test were configured to record while no input was injected into the saline bath. After obtaining 2-minute-long recordings with the above setup, root mean square (RMS) noise and the noise floor were determined. To compute noise floor values, power spectral densities (PSDs) were calculated via Welch’s method, using a 0.5s-long Hamming window and 50% overlap for the recording channel.

A segment of these raw recordings is shown in Figure 2 A-D to illustrate typical recordings from the NTBs. Despite distinct laboratory locations and setups, the NTBs showed minimal noise fluctuation across sites (Figure 2A, B, E, F). RMS values for the NTB were calculated across the entire 2-minute recordings, yielding values of 4.06µV (preclinical) and 4.01µV (clinical). After establishing consistency in the noise characteristics of the NTB between sites, the NTB was used to assess the noise floors of the amplifiers unique to each testing site. Power spectral densities were calculated for the setups at each site using activity captured from the NTB and systems under test (Figure 2E and F). The NTB showed less line noise (60Hz and harmonics) than the Intan/Open Ephys data (Figure 2E) as well as lower noise floor at all frequencies relative to the BIC (Figure 2F). Due to differences between sampling frequencies for the NTB (1-1000Hz), Intan (1-7500Hz) and BIC (1-500Hz), the PSDs for each of these recordings reached a noise floor plateau at different frequencies. The PSDs determined at the clinical site showed the BIC plateauing around 500Hz at -4dB with the NTB settling near 1000Hz at around -24dB. At the preclinical site, the PSD for the NTB also settled near 1000Hz at -24dB while the Intan setup settled near 600Hz at -25dB.

**Figure 2:**
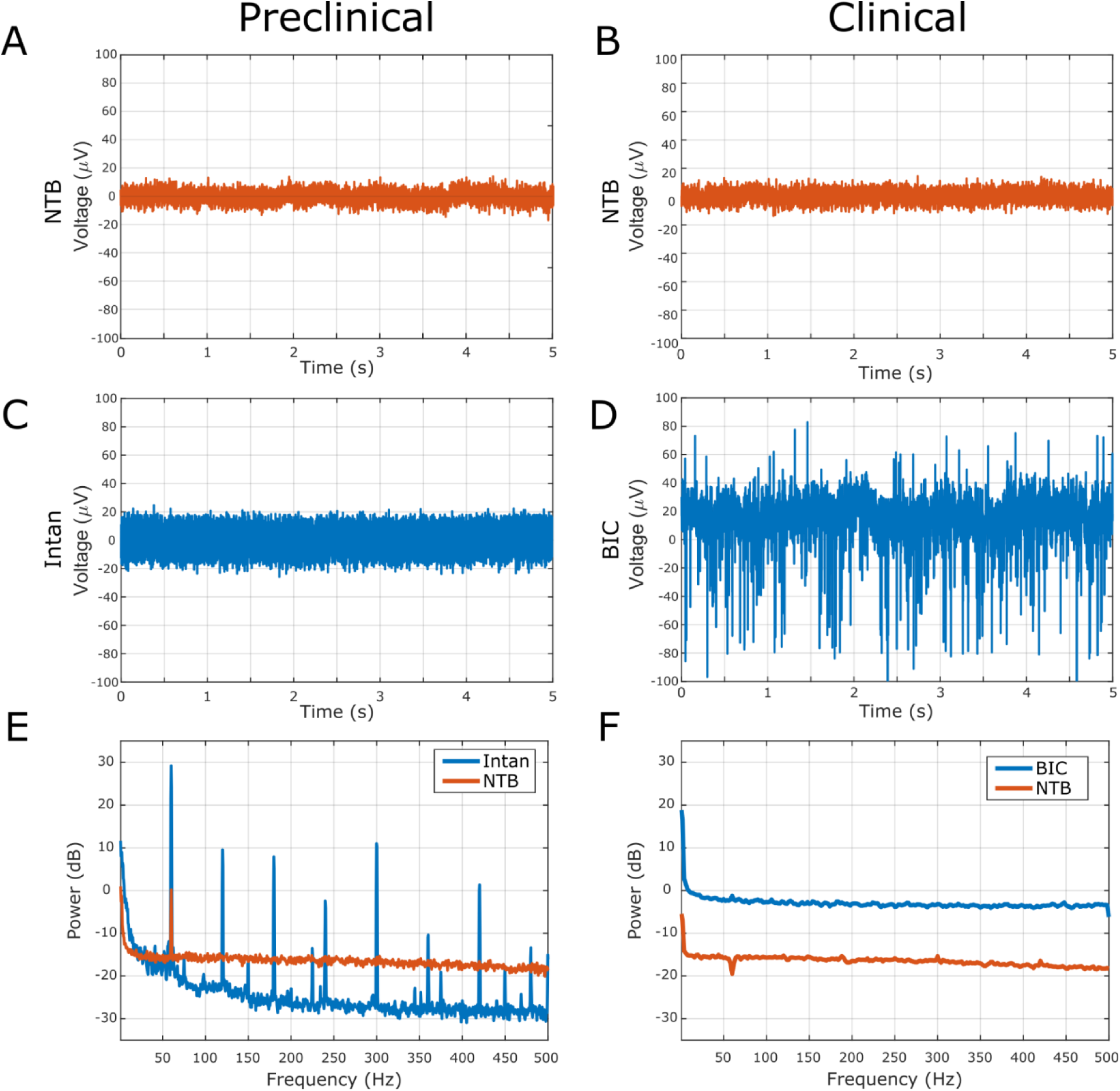
Noise floor characteristics of various data acquisition systems. Five seconds of data acquired from NTBs in saline at **(A)** the preclinical site and **(B)** the clinical site. **(C-D)** Recordings collected with an **(C)** Intan RHD2132 headstage via an Open Ephys DAQ and **(D)** CorTec BIC capturing the same activity depicted in (A) and (B), respectively. **(E)** Power spectral density plot comparing the spectral characteristics of the NTB and Open Ephys (Intan) data. While the Intan/Open Ephys data has less power at most frequencies, note the amplification of 60Hz harmonics. **(F)** Same as in (E) but comparing the NTB and the CorTec BIC. The BIC was found to have increased broad-spectrum noise relative to the NTB. With the exception of 60 Hz noise and harmonics, due to differences in filter characteristics between the sites, the two NTB systems have nearly identical spectral characteristics.

### 3.3 Evaluating Neural playback

A key functionality of the NTB is its ability to play signals of interest (e.g., previously recorded neural activity) into saline. This neural playback functionality enables a benchtop alternative to *in vivo* experimentation and, for the case of the NTB, is augmented by the system’s recording functionality. Concurrent recording of benchtop activity enables the collection of ground truth data, providing a robust means to evaluate the recording capability of a system under test.

To assess the NTB’s biopotential generation and recording capabilities, electrophysiology data collected in both preclinical and clinical contexts served as inputs to be played back by the system. Data for playback was re-sampled such that the sampling rate was 2kHz before being written to a CSV file. The CSV file was then loaded into a microSD card to be read by the NTB. Calibration was also performed prior to initiating biosignal playback. After both the NTB and systems under test captured 1min of playback (Figure 3), cross-correlation was performed to compare the similarities between the original input and captured activity by the two systems.

**Figure 3:**
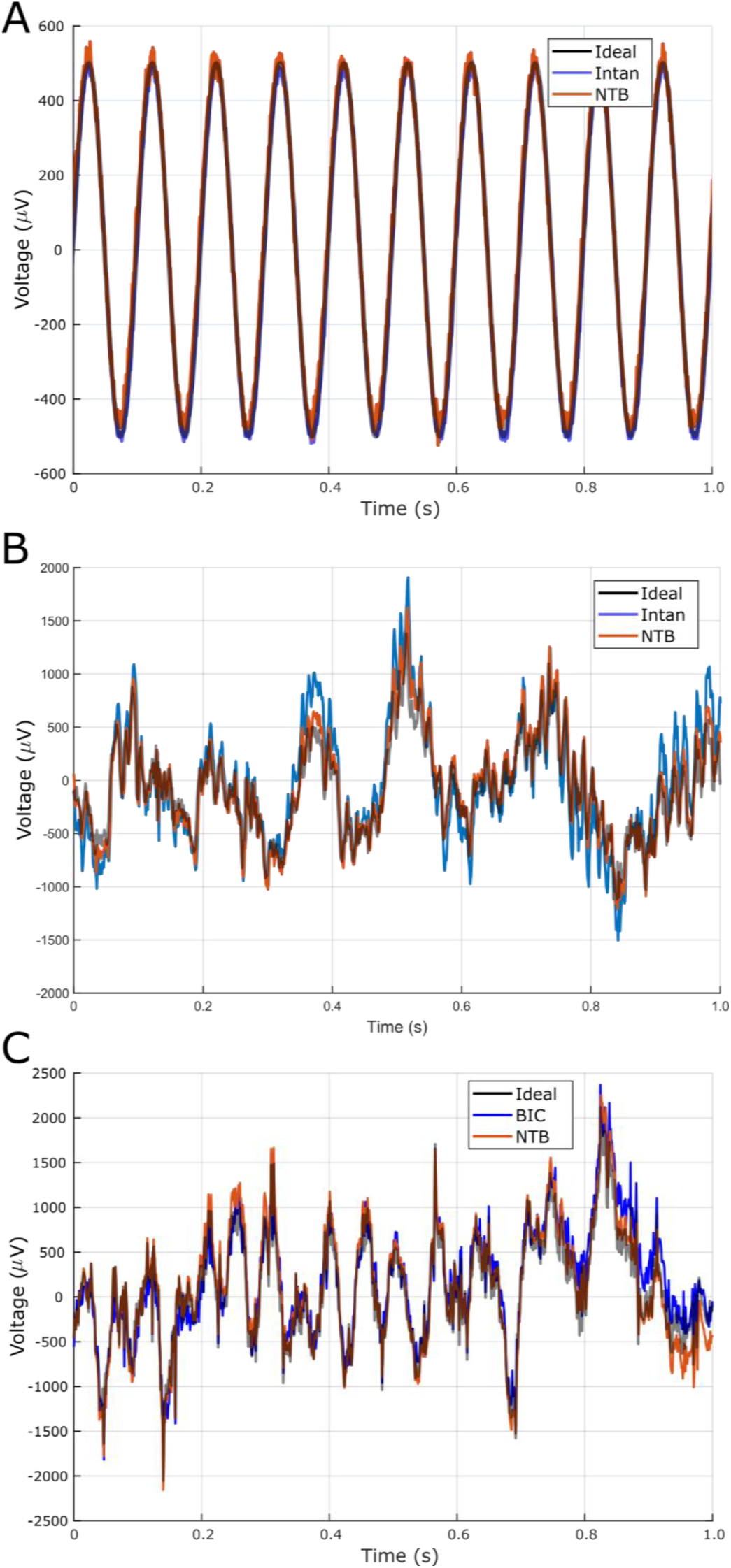
Accuracy of the current source waveform generator. **(A)** A 10Hz, 1 mVpp sine wave was injected into the saline tank for 1 s using the neural playback function. The input data (Ideal, black) is shown in black. The data recorded via the Intan RHD2132 (blue) and via the NTB (orange) is overlaid. **(B)** Neural activity collected from the rodent infralimbic cortex was injected into the preclinical site’s saline tank. One second of the recorded data is shown here comparing the ideal, Intan (Open Ephys), and NTB data streams. **(C)** Neural data collected from a patient with epilepsy was injected into the clinical site’s saline tank. One second of data comparing the Ideal, BIC (blue), and NTB (orange) is shown.

First, both sites generated CSV files containing a 10Hz, 1 mVpp sine wave and injected this signal via the neural playback function as an initial assessment of how well the ACSWG of the NTB reproduces the input signal. Figure 3A shows data collected at the preclinical site. At both sites, the correlation between the input signal and the signal collected via the NTB and devices under test was very high (preclinical: NTB r = >0.99, Intan r = >0.99; clinical: NTB r = >0.99, BIC r = 0.96).

Next, we sought to test the ACSWG with more complex signals. For the preclinical site, local field potential (LFP) data was collected from depth electrodes (Pt-Ir, 125µm diameter, <20kΩ) in the infralimbic cortex of a rat during a free-behaving recording. Animals were placed inside a non-conductive container located inside a Faraday cage and allowed to freely explore the open space for the entirety of a five-minute recording. Electrophysiology data were routed through a Brownlee pre-amplifier (Model 440) to an OpenEphys DAQ and sampled at 30kHz. These data were downsampled to 2kHz and injected into the electrophoresis tank using the neural playback function (Figure 3B). Data were routed to both the sensing portion of the NTB and to an IntanRHD2132 headstage using a splitter (described in the *Preclinical Benchtop Environment section* above). The correlation between the input signal and the NTB recording was r = 0.96, while the correlation coefficient for the input signal and Intan data was r = 0.87.

With the clinical testing environment, stereo-electroencephalography (sEEG) recordings collected during intracranial monitoring of a human participant with epilepsy were used for playback. These resting state recordings were originally collected with the CorTec BIC system connected to a subset of the implanted sEEG electrodes through externalized extensions ending in clinical touch-proof connections. From the original recordings, a select channel located in the motor cortex served as the input signal to be played back by the NTB. Signal fidelity between the input and recorded signals was again high with the correlation coefficient between the original input and the sensed activity for the NTB being r = 0.95 while the BIC registered at r = 0.86 (Figure 3C).

These data demonstrate the ability of the NTB to both generate and record complex signals while maintaining the integrity of the original dynamics of the input signal even when it is set up in non-idealized environments (e.g., mimicking a clinical space).

### 3.4 Evoked Potential Simulation

The sensing circuit is capable of communicating with the ACSWGs via the processing unit, which enables experimenters to generate reactions to specific stimuli. When the sensing circuit detects an event of interest, the processing unit can trigger the ACSWG to produce a pre-programmed response. While the user can set the specific parameters that define a triggering event, we initially programmed the NTB to detect DBS-like stimulation pulses (i.e., large changes in energy at high frequencies) and react with a simulated evoked potential that is defined by four customizable parameters to emulate an evoked response expected *in vivo*.

(1) The default amplitude range of the output is 1µV to 2000µV (Figure 4A). The maximum of 2000µV can be overridden by the end user, however, the maximum voltage range will be limited by the test setup maximum impedance as the ACSWG can generate currents up to 10mA with a maximum voltage range of ±14V. Note, achieving an accurate voltage range requires use of the calibration function prior to EP testing.
(2) The latency of the EP following a detected event of interest (e.g., the stimulation artifact from an electrical pulse) can range from 0 to 1s (Figure 4B). The upper limit of this delay parameter can be programmatically adjusted.
(3) The frequency of the EP can be set between 1Hz and 250Hz (Figure 4C). The cap of the frequency component can be increased programmatically by increasing the write speed of the DAC and changing the hardcoded frequency maximum.
(4) The final adjustable feature of the EP is the “template” of the response. The NTB software currently supports three different EP templates to account for various EP morphologies (Figure 4D): **Type I** features one full cycle of a sine wave at a single frequency in which the first half of the cycle is delivered at 100% amplitude and the second half of the cycle is delivered at 25% amplitude. The frequency and amplitude of the EP are set by the user. The sine wave has a negative leading phase, resulting in a starting voltage near 0 followed by a negative trough (Figure 4D, top). **Type II** consists of a cosine wave with a negative leading phase and the same amplitude decay as Type I (i.e., 100% for the first half cycle and 25% for the second half). This results in a positive voltage at EP onset that falls to a trough before moving to a lower amplitude positive peak (Figure 4D, middle). Finally, **Type III** features the combination of two sine waves at different frequencies set by the user. The first sine wave is delivered for one half cycle at 100% amplitude, while the second is delivered for a full cycle, with the first half at 100% amplitude the second half at 25% amplitude (Figure 4D, bottom).

**Figure 4:**
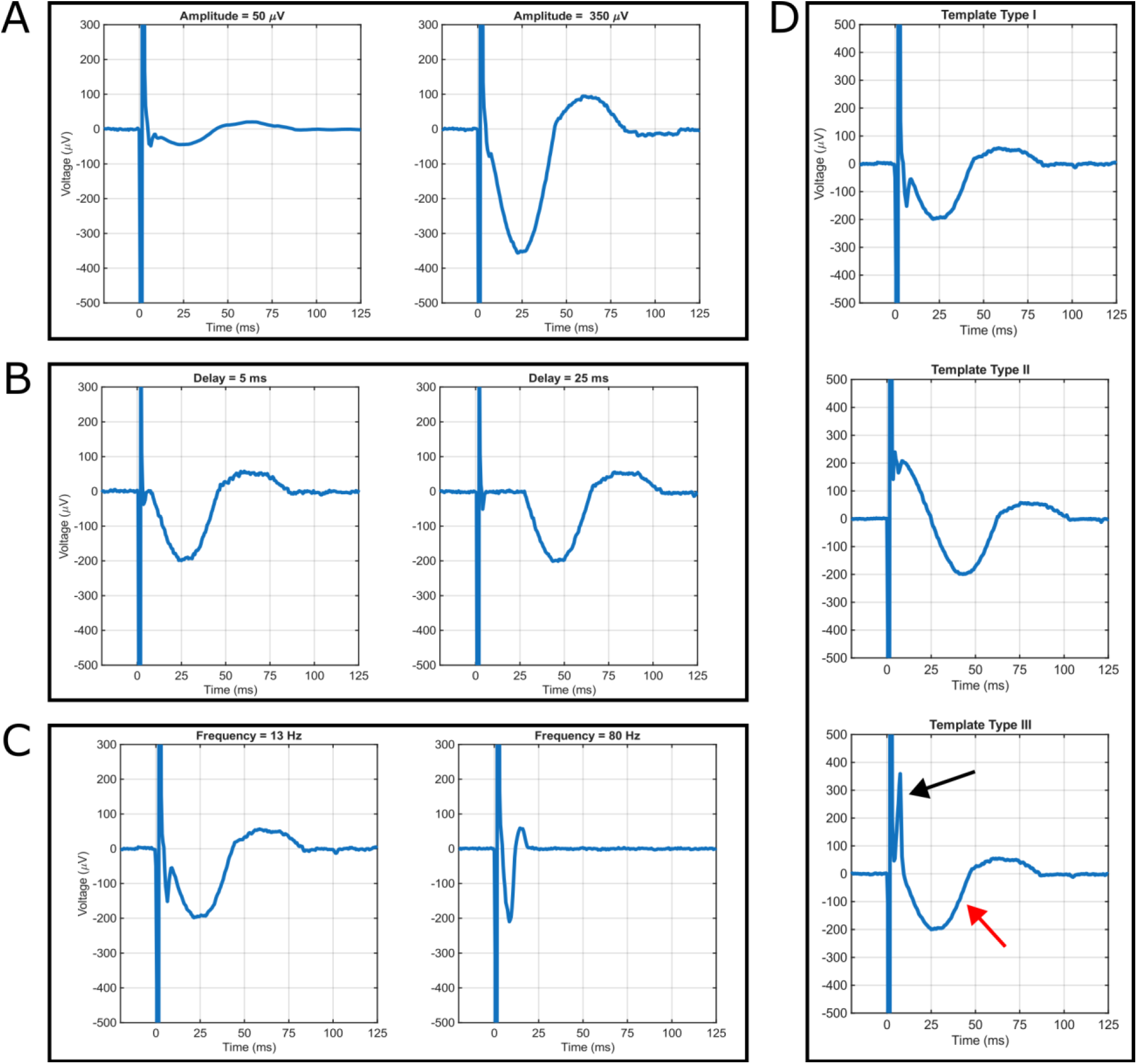
Configurable parameters for evoked potential simulation with the NTB. **(A)** Example of changing the amplitude component of the stimulated evoked potential (EP). Both EPs used a 13Hz sine wave as a base. The left panel shows a 50µV EP, while the right shows a 350µV EP. **(B)** Example of changing the delay from stimulation detection to EP generation. Both EPs used a 13Hz, 200µV amplitude sine wave as a base. The left side shows an EP delayed by 5ms while the right shows a 25ms delay. **(C)** Example of changing the frequency component of the EP. Both used a 200µV sine wave as a base. The left side shows an EP generated from a 13Hz sine wave while the right shows an EP generated from an 80Hz sine wave. **(D)** Example of changing the morphology of the evoked potential between three configurations. Type I - a negative leading sine wave in which the first half of the cycle is delivered at 100% amplitude, and the next half cycle is delivered at 25% amplitude. Type II - a single frequency, negative leading cosine wave where the first oscillation is delivered at 100% amplitude and the second half cycle is delivered at 25% amplitude. Type III - a leading high frequency, negative half sine wave (black arrow) followed by a low frequency full sine wave (red arrow) where the ratio between the two frequencies is 20 and the ratio between the two amplitudes is 2. The high frequency component is 100 Hz in this example.

In this stimulation context, the NTB’s data acquisition capability is particularly relevant, as stimulation can overwhelm a system’s amplifier, resulting in gaps of information with respect to the impact of stimulation on capturing evoked neural activity. By having ground-truth knowledge of the actual “biosignal” in saline, the effects of stimulation and its artifacts can be characterized and not only inform the design of stimulation experiments but also guide signal processing methods for post-hoc analysis. Considering the variety of stimulation evoked activity, the NTB’s EP simulation functionality allows customization of the evoked response, providing an initial starting point for modeling biologically plausible evoked potentials. We describe tests using this functionality below.

## 4. Applications of the NeuroTest platform

### 4.1 Evaluating recording configurations for evoked potential experiments

We used the NTB to evaluate whether a simulated EP could be recovered from data collected via an Intan RHD2132 headstage. These headstages are common in preclinical electrophysiology applications but were not designed with electrical stimulation in mind (see the Intan RHS series for devices designed around stimulation experimentation). Despite this, several groups have reported electrical stimulation experiments using these chips with varying degrees of success at artifact mitigation (see [40–42] for a subset). This makes the RHD2132 chip an interesting edge-case: a device that appears capable of usage in stimulation experiments under certain conditions but one that lacks standardized or manufacturer testing within this specific context.

To assess how quickly a known signal could be recovered following stimulation when using an IntanRHD2132, we used an external stimulator (StimJim) to deliver biphasic stimulation pulses (+/-100µA, 100µs per phase) into the electrophoresis tank via a pair of electrodes (∼4kΩ impedances). This pulse was detected by the sensing portion of the NTB and a parallel data stream routed through an Intan RHD2132 to an Open Ephys data acquisition system. The simulated EP used the Template Type I format (i.e., produces one cycle of a 25Hz sinewave where the first half of the wave is delivered at 100% amplitude and the next half at 25% amplitude). The sine wave was set to have a target realized amplitude of 150µV. In addition to being within a reasonable amplitude range of reported CCEP components [43,44], this value was selected in order to ensure a detectable response that could also allow thorough assessment of the efficacy of post-hoc processing techniques to extract the underlying EP activity. Stimulation in the Intan context resulted in a large post-stimulation artifact that required ∼600ms to settle back to baseline (Figure 5A and B). To isolate the simulated EP, we employed a template subtraction technique. We found that the stimulation artifact present in the Intan data stream was not consistent (as has been the case in our *in vivo* data, not shown), making subtraction of a single template challenging. Instead, we created a series of artifact templates by delivering 20 stimulation pulses via the StimJim while the evoked potential function of the NTB was disabled. We then delivered sets of 5 stimulation pulses with the evoked potential function active using either a 1ms delay (Figure 5A, C, and E) or 25ms delay (Figure 5B, D, and F). We time-aligned each of the 5 evoked potential pulses with each of the 20 template artifacts and did a timepoint-by-timepoint subtraction of the templates from the evoked potential pulses. We then squared and summed the resulting values and selected the pairing with the lowest sum of squares as the best fit template (shown in Figure 5C and D). We further verified accuracy of the template by calculating the correlation coefficient between the artifact template and evoked potential pulse (Figure 5C and D). In all cases, the correlation coefficients were r = >0.99. Finally, we take the result of subtracting the best fit template from the raw data to show that the evoked potential was recoverable (Figure 5E and F) despite the large, inconsistent post-stimulation artifact present in the Intan data stream. Development of a similar template matching strategy for *in vivo* data would be challenging as the known ground truth for the EP would not be available. However, this test verifies that the RHD2132 data stream does contain the EP embedded within the post-stimulation artifact, enabling future attempts at *in vivo* EP recovery.

**Figure 5:**
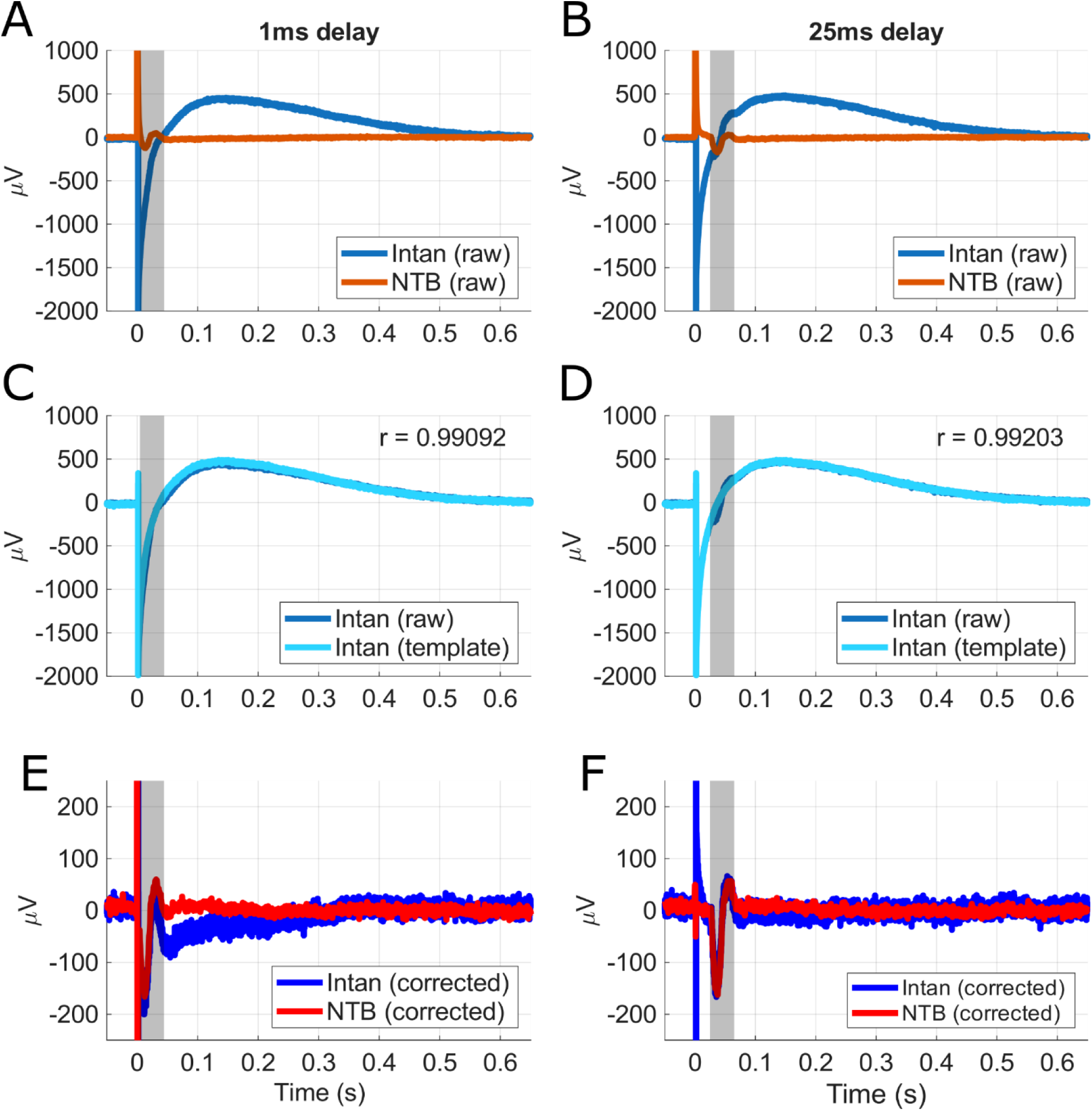
Using the NTB to develop an artifact mitigation approach based on simple template matching. **(A)** Raw time series data from the NTB (orange) and Intan RHD2132 via Open Ephys (dark blue) setups depicting an example stimulation pulse and simulated EP. Gray shaded area shows the expected start and end times of the simulated EP (parameters: 25Hz sine wave, 150µV target amplitude, Template Type 1, 1ms delayed response relative to detection of stimulation pulse). Note the large post-stimulation artifact in the Intan data stream and the lack of a visually detectable EP. **(B)** Same as in A but using a 25ms delay for the EP generation. **(C)** Same data from the Intan stream as shown in A but with a template matched artifact overlaid (light blue). The correlation coefficient between the artifact template and the raw artifact is displayed. **(D)** Same as in C but using a 25ms delay. **(E)** Simulated EPs extracted from the NTB (red) and the Intan (blue) data streams using a template subtraction method. **(F)** Same as in E but using a 25ms delayed EP. Note that the voltage scale has been changed from A-D to visualize the precision of the amplitude of the EP following artifact rejection.

### 4.2 Testbed for evoked potential processing

We utilized the NTB’s EP simulation feature to assess the ability of the BIC to reliably capture stimulation evoked activity and evaluate the efficacy of post-hoc cleaning techniques for isolating the EP of interest collected via the BIC system. Using the recording configuration described previously, low-frequency stimulation pulses were delivered by the BIC and triggered a consistent evoked response from the NTB. With the command interface, a 150µVpp signal with 12Hz frequency components was injected immediately upon detecting stimulation. This evoked response contained an initial negative peak around 25ms post-stimulation, and a smaller positive peak about 60ms post-stimulation. Throughout these stimulation experiments, both the BIC and NTB systems were set to record. For this characterization, 20 trials of stimulation were delivered with the BIC. During post-hoc analysis, the trials were epoched and then averaged to generate an average of the raw evoked response waveform.

Trial data and their raw averages from this stimulation experiment can be seen for the BIC and NTB systems in Figure 6A and B, respectively. The raw average computed from the BIC recordings shows the evoked response signal superimposed with an exponential trend as the amplifier recovered from stimulation. The much larger magnitude of the stimulation artifact and low-frequency nature of the return to baseline behavior made it particularly challenging to fully appreciate the underlying evoked response. With further inspection of individual trial outcomes, return to baseline behavior, particularly activity within 150ms post stimulation, was highly variable, distorting the underlying evoked potential response. Activity captured by the NTB shows the stimulation artifact generated by the BIC and the preserved components of the generated evoked response. Upon comparison between the NTB and BIC trials and averages, the variable exponential decay from stimulation was unique to the BIC dataset only, indicating that this phenomenon is specific to the BIC device.

**Figure 6:**
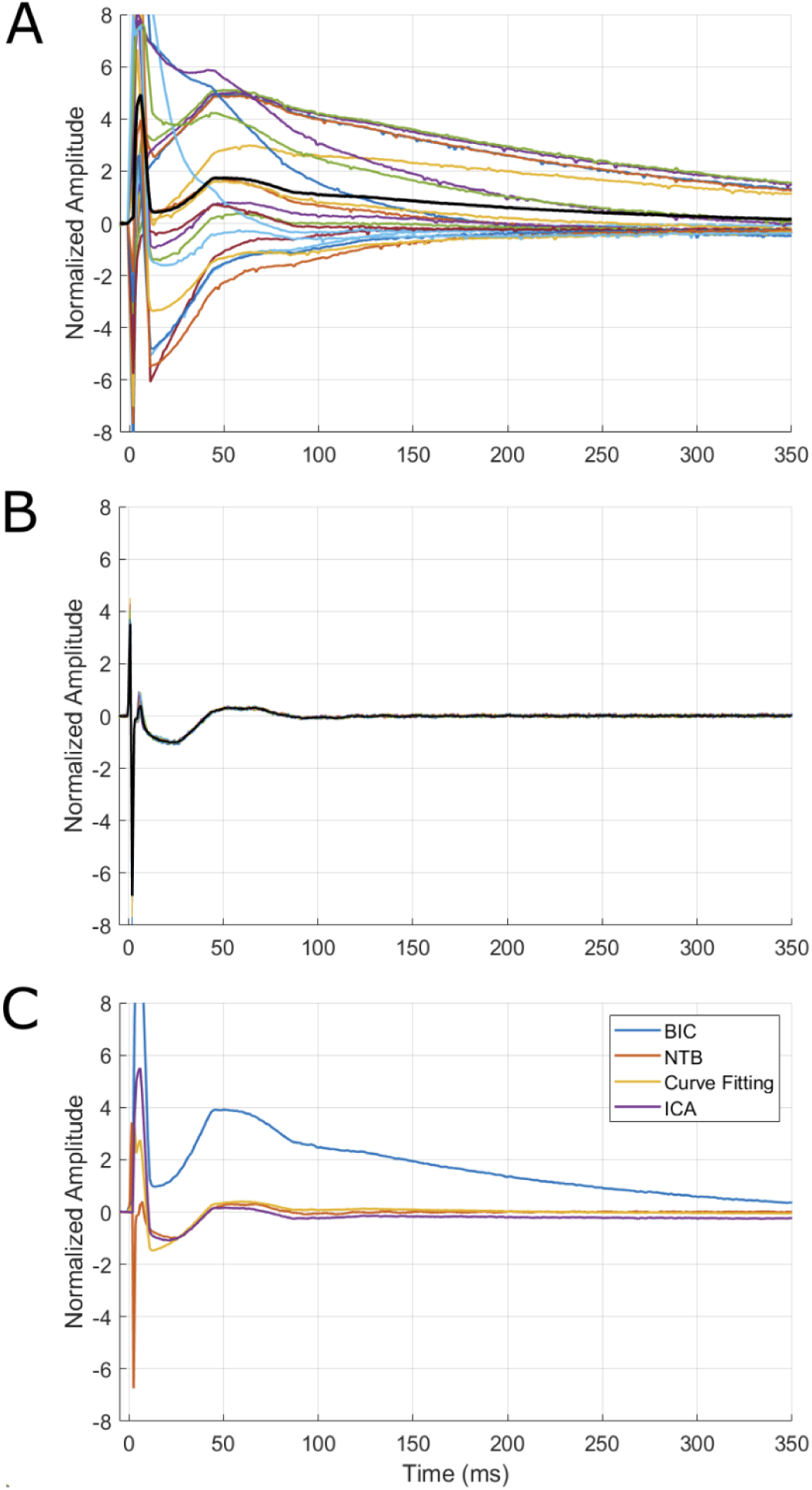
Example case of using the NTB and its evoked potential functionality to assess signal processing methods. Time synchronized plots of normalized, simulated evoked potentials (parameters: 12Hz, 150µV, Template Type 1, no delay) as captured by **(A)** the BIC and **(B)** the NTB. Colored traces show individual trial data while the raw average of the trials is plotted in black. **(C)** Outcomes from applying ICA and curve fitting approaches to target the return to baseline behavior are illustrated and compared to the ground truth recorded by the NTB and the original average response from the BIC. Both signal processing approaches targeted the low-frequency return to baseline observed in the original BIC trace. Normalized amplitudes facilitated comparison of cleaned responses between signal processing techniques.

Because of interest in the temporal characteristics of evoked activity, we implemented different strategies to minimize the impact of the exponential trend on the evoked response. In this application, we applied a curve-fitting approach and independent component analysis (ICA) to clean the BIC dataset. To facilitate comparisons between these methods, signal outcomes from these cleaning methods, Y_C_, were normalized. For assessing the efficacy of these methods to extract the underlying evoked responses, the NTB outcomes, Y_O_, were used as an effective ground truth dataset to compute the relative error of the cleaned signals. Relative error was calculated with the following equation where || x ||_2_ denotes the Euclidean norm of the signal, x:

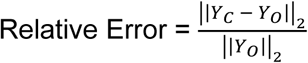

We first utilized the curve-fitting approach. The average response from utilizing this method can be seen in Figure 6C. To target the exponential trends captured during these stimulation experiments, an exponential curve was modeled to the overall trend of the trial before being subtracted from the trial data. This method towards detrending was somewhat effective towards better realizing the evoked response. While this approach helped offset the exponential trend, it did not fully recover all components of the simulated response, in particular, activity that occurred during or shortly after stimulation. Additionally, it was noted that poor fits can result in the creation of pseudo-oscillations that could be mistakenly interpreted as electrophysiological activity or distort the underlying oscillations of interest. Considering that the underlying EP is an unknown signal, an additional vetting process should be incorporated to ensure the quality of a fit prior to detrending. However, it should be noted that while metrics such as root mean square error could potentially mitigate the application of poor fits to detrend data, it may be difficult to determine the threshold to discern a poor fit due to the presence of underlying activity. In using this curve-fitting approach, the relative error from utilizing this method was computed to be 50%.

Another method applied was independent component analysis (ICA). Prior efforts have utilized ICA for targeting artifacts and their distortions in various stimulation evoked response experiments [45,46]. Using the FastICA method [47], ICA components were selectively removed to eliminate the stimulation artifact recovery period overwhelming the underlying evoked potential response. Because of the low-frequency nature of the return to baseline behavior and possible overlap with the lower frequency nature of the simulated signal, components were manually identified prior to removal. This approach resulted in reasonable recovery of the components of the simulated response. Compared to the curve-fitting outcomes, application of ICA allowed the recovery of the negative peak 25ms post-stimulation. However, it should be noted that this approach must be customized to each dataset, as evoked activity can be composed of various neural sources. With ICA, the relative error was determined to be 36%. Outcomes with the ICA approach resulted in slightly improved recovery compared to the curve-fitting approach; however, evaluation of additional processing methods is of interest in order to yield further improvements in signal recovery. While a thorough investigation of artifact mitigation techniques is beyond the scope of this work, these results demonstrate that the NTB can be used to facilitate these tests.

## 5. Discussion

In this work, we present the NTB, a controlled, benchtop evaluation system that enables extensive device characterization and validation to guide neuromodulation research efforts. The NTB and electrophoresis tank provide a platform that is designed to mimic bio-electric signals as well as the tissue-electrode interface, allowing for *in vitro* alternatives to *in vivo* testing. Assessments across labs demonstrated highly consistent performance of the NTB despite distinct environments, including differences in the devices (i.e., Intan RHD2132 and CorTec BIC). Noise floor characteristics and signal generation fidelity of the NTBs were nearly identical, establishing a reliable benchmark for assessing functionality of other devices. We further provide two separate use-cases of the neural playback functionality to assess stimulation artifacts in simulated EP experiments. Taken together, these data demonstrate that the NTB can be used for critical testing and validation of brain stimulation and sensing technologies across sites, devices, and applications.

Reproducibility is critical for scientific research [48–51] and the need for reliable benchmarks will only increase as electrophysiology experiments and related technologies become more complex. This reality is the core motivation for the creation of open-source devices like the NeuroDAC [22] and our NTB. Having systems that can be used flexibly across many different applications and experimental setups allows for comparison across projects with disparate methodologies but overlapping scientific purposes. In our case, the teams at each site have similar end goals (i.e., developing new neuromodulation therapies for neurological and neuropsychiatric disorders); however, testing at the first site was done in a preclinical setting while testing second site was done in a clinical laboratory environment. The preclinical team used electrodes, stimulators, and data acquisition systems common in animal model-based research while the clinical team worked with a neuromodulation device in development for human use. We demonstrate that the NTB behaves predictably and similarly across these two laboratory setups despite their different methodological approaches.

The noise floors on our NTB devices follow a highly consistent and straightforward power-law scaling that starts below 0dB and bottoms out around -25dB. General background noise in both systems was ∼4µV RMS despite the lack of faraday cages or other electrical isolation and environmental noise mitigation techniques often present in electrophysiology rigs. Given that neural signals recorded via intracranial electrodes generally have amplitudes in the 1-500µV range, these data show that the NTB can reliably detect even small simulated biopotentials. We further demonstrate that the signal generator of the NTB exhibits extremely high-fidelity replay capabilities both with simple signals (i.e., sine waves) and complex neural data from both rodents and humans. These data highlight the potential for this system to be used across the translational pipeline, from preclinical animal modeling to evaluation of investigational devices.

A core use case of the NTB is emulating evoked potentials. This functionality allows us to conduct experiments assessing the impact of stimulation artifacts and device amplifier limitations on known input signals (i.e., the simulated EPs). We used this functionality to show that the commonly used Intan RHD2132 digital headstage could reliably be used to record simulated EPs within 1ms of a large stimulation artifact under our specific testing conditions, but only with a known artifact template that allowed subtraction. The large amplitude and rapid exponential/non-linear decay of the post-stimulation artifact makes assessing any underlying physiology difficult, particularly in an *in vivo* context where a neural evoked response may or may not be present. The inconsistency of the post-stimulation artifacts further complicates any *in vivo* assessment. Here, we leveraged two main components to assess signal fidelity: (1) the parallel data stream routed to the NTB, which remains largely free from post-stimulation artifacts and (2) the known ground truth of the simulated evoked potential. We provide a simple methodology for artifact removal in benchtop testing that can be validated against these two components. This approach allows users to find the necessary conditions to recover neural data post-stimulation as rapidly as possible without the need for complicated and uncertain *in vivo* testing. We want to again note that we used the RHD2132 due to its ubiquity and because our NTB testing application represented an edge case for use of this equipment. The Intan RHS series is an alternative system purpose-built for electrical stimulation experiments.

In our clinical testing space, we were able to assess the post-stimulation artifacts observed with the CorTec BIC. With a lower sampling frequency and more convoluted return to baseline behavior, the BIC system requires more advanced post-hoc techniques to extract the underlying response of interest. Simulations of CCEP experiments with the NTB allowed us to generate a data set to evaluate analytical techniques for artifact removal. In both sets of tests, outcomes from the cleaning methods were compared to the known generated outputs captured by the NTB. Assessments with this ground truth dataset identified nuances in these processing methods and set up an initial pipeline that can be applied when approaching *in vivo* CCEP collection.

Although the presented NeuroTest platform expands on current biopotential emulation systems, there are notable limitations with the NTB system. Currently, the NTB only supports two input and output channels, placing a limit on the complexity that can be emulated in this testing environment. This configuration may not be appropriate for emulating more complex environments, particularly within applications exploring multi-source brain activity (e.g., cross frequency coupling or coherence [52,53]) or investigating spatio-temporal sensing configurations (e.g., travelling waves [54] or topographical functional mapping [55–57]) [4,58,59]. Expansion of the current circuitry is relatively straightforward, though, and would mainly require duplication of the existing architecture. Another shortcoming of the present NTB is the use of a microSD to load data available for playback. While advances in memory technology have enabled significant increases in storage capacity, the process of modifying data for playback, reuploading the data to the memory card, and then physically mounting the card to the NTB can be laborious and serve as an inconvenient barrier for rapid iteration through various signals for testing. One alternative is to allow direct signal loading via serial commands, improving flexibility with the device that can be enabled with firmware and software development. Additionally, while the NTB can support a range of signals for playback, emulation of spiking activity is limited by the settling time of the DAC which is 1μS. Likewise, the ADC sampling rate is set to 1kHz by default, meaning that this device is currently optimized for applications involving meso-to macro-scale neural activity. However, increasing the maximum speed of the signal generator is possible through removal of the feedback capacitor, while the ADC sampling rate can be programmatically set to higher frequencies (up to 32kHz). These adjustments could be made to enable the high temporal precision needed for fine grained emulation of action potentials, though further testing would be warranted.

In the future, the neural playback and EP functionalities of the NTB can be used for more sophisticated assessments of closed-loop algorithms and experimental designs. The open-source code could be updated to allow for stimulation parameters that adapt over time, or the system could be programmed to cycle through evolving simulated neural states to mimic neuroplastic changes over time. We intend to continue to improve the NTB by addressing the hardware and software limitations noted above in order to facilitate experiments that feature embedded neuronal models, multi-source/network level interactions, and optimization of stimulation parameters in implanted neuromodulatory systems [24,60–62].

## 6. Conclusion

With advances in the neuromodulation space, the need for evaluation tools will play a significant role in ensuring adequate preparation before starting *in vivo* research. In building off of currently available evaluation systems, there will be more accessible means to assess electrophysiology equipment and experimental setups. Better informing users of the nuances of the systems they are utilizing in novel research efforts will guide upcoming experiment design and facilitate improved hardware design for future equipment. By expanding the resources available to perform system evaluation work, researchers can raise confidence in the novel systems they are utilizing and support more rapid and reliable prototyping efforts within the neural interface and neuromodulation space.

## REFERENCES

[1] Konrad P, Gelman K R, Lawrence J, Bhatia S, Jacqueline D, Sharma R, Ho E, Byun Y W, Mermel C H and Rapoport B I 2025 First-in-human experience performing high-resolution cortical mapping using a novel microelectrode array containing 1024 electrodes *J*. Neural Eng. 22

[2] Kullmann A, Kridner D, Mertens S, Christianson M, Rosa D and Diaz-Botia C A 2022 First Food and Drug Administration Cleared Thin-Film Electrode for Intracranial Stimulation, Recording, and Monitoring of Brain Activity-Part 1: Biocompatibility Testing *Front*. Neurosci. 16 876877

[3] Trautmann E M, Hesse J K, Stine G M, Xia R, Zhu S, O’Shea D J, Karsh B, Colonell J, Lanfranchi F F, Vyas S, Zimnik A, Amematsro E, Steinemann N A, Wagenaar D A, Pachitariu M, Andrei A, Lopez C M, O’Callaghan J, Putzeys J, Raducanu B C, Welkenhuysen M, Churchland M, Moore T, Shadlen M, Shenoy K, Tsao D, Dutta B and Harris T 2025 Large-scale high-density brain-wide neural recording in nonhuman primates *Nat*. Neurosci. 28 1562–75

[4] Herron J, Kullmann A, Denison T, Goodman W K, Gunduz A, Neumann W-J, Provenza N R, Shanechi M M, Sheth S A, Starr P A and Widge A S 2025 Challenges and opportunities of acquiring cortical recordings for chronic adaptive deep brain stimulation *Nat*. Biomed. Eng. 9 606–17

[5] Jeong J-W, Shin G, Park S I, Yu K J, Xu L and Rogers J A 2015 Soft Materials in Neuroengineering for Hard Problems in Neuroscience Neuron 86 175–86

[6] Widge A S 2024 Closing the loop in psychiatric deep brain stimulation: physiology, psychometrics, and plasticity Neuropsychopharmacology 49 138–49

[7] Bronte-Stewart H M, Beudel M, Ostrem J L, Little S, Almeida L, Ramirez-Zamora A, Fasano A, Hassell T, Mitchell K T, Moro E, Gostkowski M, Chattree G, de Bie R M A, de Neeling M, Piña-Fuentes D, Swinnen B, Starr P A, Hammer L H, Foote K D, Richardson R M, Flaherty A, Boogers A, Sa’di Q, Meoni S, Castrioto A, Stanslaski S, Summers R L S, Tonder L, Tan Y, Berrier H, Goble T J, Raike R S, Herrington T M, and ADAPT-PD Investigators 2025 Long-Term Personalized Adaptive Deep Brain Stimulation in Parkinson Disease: A Nonrandomized Clinical Trial JAMA Neurol.

[8] Heck C N, King-Stephens D, Massey A D, Nair D R, Jobst B C, Barkley G L, Salanova V, Cole A J, Smith M C, Gwinn R P, Skidmore C, Van Ness P C, Bergey G K, Park Y D, Miller I, Geller E, Rutecki P A, Zimmerman R, Spencer D C, Goldman A, Edwards J C, Leiphart J W, Wharen R E, Fessler J, Fountain N B, Worrell G A, Gross R E, Eisenschenk S, Duckrow R B, Hirsch L J, Bazil C, O’Donovan C A, Sun F T, Courtney T A, Seale C G and Morrell M J 2014 Two-year seizure reduction in adults with medically intractable partial onset epilepsy treated with responsive neurostimulation: Final results of the RNS System Pivotal trial Epilepsia 55 432–41

[9] Scangos K W, Khambhati A N, Daly P M, Makhoul G S, Sugrue L P, Zamanian H, Liu T X, Rao V R, Sellers K K, Dawes H E, Starr P A, Krystal A D and Chang E F 2021 Closed-loop neuromodulation in an individual with treatment-resistant depression *Nat*. Med. 27 1696–700

[10] Figee M, Riva-Posse P, Choi K S, Bederson L, Mayberg H S and Kopell B H 2022 Deep Brain Stimulation for Depression Neurotherapeutics 19 1229–45

[11] Sheth S A, Bijanki K R, Metzger B, Allawala A, Pirtle V, Adkinson J A, Myers J, Mathura R K, Oswalt D, Tsolaki E, Xiao J, Noecker A, Strutt A M, Cohn J F, McIntyre C C, Mathew S J, Borton D, Goodman W and Pouratian N 2022 Deep Brain Stimulation for Depression Informed by Intracranial Recordings *Biol*. Psychiatry 92 246–51

[12] Ansó J, Benjaber M, Parks B, Parker S, Oehrn C R, Petrucci M, Gilron R, Little S, Wilt R, Bronte-Stewart H, Gunduz A, Borton D, Starr P A and Denison T 2022 Concurrent stimulation and sensing in bi-directional brain interfaces: a multi-site translational experience *J*. Neural Eng. 19 026025

[13] Stanslaski S, Herron J, Chouinard T, Bourget D, Isaacson B, Kremen V, Opri E, Drew W, Brinkmann B H, Gunduz A, Adamski T, Worrell G A and Denison T 2018 A Chronically-Implantable Neural Coprocessor for Investigating the Treatment of Neurological Disorders. IEEE Trans. Biomed. Circuits Syst. 12 1230–45

[14] Elder C M, Hashimoto T, Zhang J and Vitek J L 2005 Chronic implantation of deep brain stimulation leads in animal models of neurological disorders *J. Neurosci*. Methods 142 11–6

[15] Baker J L, Ryou J-W, Wei X F, Butson C R, Schiff N D and Purpura K P 2016 Robust modulation of arousal regulation, performance, and frontostriatal activity through central thalamic deep brain stimulation in healthy nonhuman primates *J*. Neurophysiol. 116 2383–404

[16] Dastin-van Rijn E M, Provenza N R, Calvert J S, Gilron R, Allawala A B, Darie R, Syed S, Matteson E, Vogt G S, Avendano-Ortega M, Vasquez A C, Ramakrishnan N, Oswalt D N, Bijanki K R, Wilt R, Starr P A, Sheth S A, Goodman W K, Harrison M T and Borton D A 2021 Uncovering biomarkers during therapeutic neuromodulation with PARRM: Period-based Artifact Reconstruction and Removal Method *Cell Rep*. Methods 1 100010

[17] Opri E, Isbaine F, Borgheai S B, Bence E, Deligani R J, Willie J T, Gross R E, Au Yong N and Miocinovic S 2025 Deep brain stimulation-induced local evoked potentials outperform spectral features in spatial and clinical STN mapping *J*. Neural Eng. 22 046055

[18] Hammer L H, Kochanski R B, Starr P A and Little S 2022 Artifact characterization and a multipurpose template-based offline removal solution for a sensing-enabled deep brain stimulation device *Stereotact*. Funct. Neurosurg. 100 168–83

[19] Chen P, Kim T, Dastin-van Rijn E, Provenza N R, Sheth S A, Goodman W K, Borton D A, Harrison M T and Darbon J 2022 Periodic Artifact Removal With Applications to Deep Brain Stimulation *IEEE Trans*. Neural Syst. Rehabil. Eng. Publ. IEEE Eng. Med. Biol. Soc. 30 2692–9

[20] Blackrock Neurotech 2015 Digital Neural Signal Simulator Blackrock Neurotech

[21] Plexon 2021 Headstage Tester Units Plexon

[22] Powell M P, Anso J, Gilron R, Provenza N R, Allawala A B, Sliva D D, Bijanki K R, Oswalt D, Adkinson J, Pouratian N, Sheth S A, Goodman W K, Jones S R, Starr P A and Borton D A 2021 NeuroDAC: an open-source arbitrary biosignal waveform generator *J*. Neural Eng. 18 016010

[23] Haci D, Liu Y and Constandinou T G 2017 32-Channel ultra-low-noise arbitrary signal generation platform for biopotential emulation 2017 IEEE International Symposium on Circuits and Systems (ISCAS) 2017 IEEE International Symposium on Circuits and Systems (ISCAS) pp 1–4

[24] Stanslaski S, Farooqi H, Sanabria D E and Netoff T I 2022 Fully Closed Loop Test Environment for Adaptive Implantable Neural Stimulators Using Computational Models *J*. Med. Devices 16

[25] International Organization for Standardization 2017 ISO 14708-3 Implants for surgery — Active implantable medical devices Part 3: Implantable neurostimulators

[26] Miller K J, Müller K-R, Valencia G O, Huang H, Gregg N M, Worrell G A and Hermes D 2023 Canonical Response Parameterization: Quantifying the structure of responses to single-pulse intracranial electrical brain stimulation *PLOS Comput*. Biol. 19 e1011105

[27] Crocker B, Ostrowski L, Williams Z M, Dougherty D D, Eskandar E N, Widge A S, Chu C J, Cash S S and Paulk A C 2021 Local and distant responses to single pulse electrical stimulation reflect different forms of connectivity NeuroImage 237 118094

[28] Paulk A C, Zelmann R, Crocker B, Widge A S, Dougherty D D, Eskandar E N, Weisholtz D S, Richardson R M, Cosgrove G R, Williams Z M and Cash S S 2022 Local and distant cortical responses to single pulse intracranial stimulation in the human brain are differentially modulated by specific stimulation parameters Brain Stimulat. 15 491–508

[29] Mekhail N A, Levy R M, Deer T R, Kapural L, Li S, Amirdelfan K, Pope J E, Hunter C W, Rosen S M, Costandi S J, Falowski S M, Burgher A H, Gilmore C A, Qureshi F A, Staats P S, Scowcroft J, McJunkin T, Carlson J, Kim C K, Yang M I, Stauss T, Petersen E A, Hagedorn J M, Rauck R, Kallewaard J W, Baranidharan G, Taylor R S, Poree L, Brounstein D, Duarte R V, Gmel G E, Gorman R, Gould I, Hanson E, Karantonis D M, Khurram A, Leitner A, Mugan D, Obradovic M, Ouyang Z, Parker J, Single P and Soliday N 2024 ECAP-controlled closed-loop versus open-loop SCS for the treatment of chronic pain: 36-month results of the EVOKE blinded randomized clinical trial *Reg*. Anesth. Pain Med. 49 346–54

[30] Keller C J, Huang Y, Herrero J L, Fini M E, Du V, Lado F A, Honey C J and Mehta A D 2018 Induction and Quantification of Excitability Changes in Human Cortical Networks *J*. Neurosci. 38 5384–98

[31] Levinson L H, Sun S, Paschall C J, Perks K M, Weaver K E, Perlmutter S I, Ko A L, Ojemann J G and Herron J A 2024 Data processing techniques impact quantification of cortico-cortical evoked potentials *J. Neurosci*. Methods 408 110130

[32] Texas Instruments Incorporated 2013 AN-1515 A Comprehensive Study of the Howland Current Pump

[33] Buzsáki G, Anastassiou C A and Koch C 2012 The origin of extracellular fields and currents — EEG, ECoG, LFP and spikes *Nat*. Rev. Neurosci. 13 407–20

[34] Goyal A, Goetz S, Stanslaski S, Oh Y, Rusheen A E, Klassen B, Miller K, Blaha C D, Bennet K E and Lee K 2021 The development of an implantable deep brain stimulation device with simultaneous chronic electrophysiological recording and stimulation in humans *Biosens*. Bioelectron. 176 112888

[35] Olsen S T, Basu I, Bilge M T, Kanabar A, Boggess M J, Rockhill A P, Gosai A K, Hahn E, Peled N, Ennis M, Shiff I, Fairbank-Haynes K, Salvi J D, Cusin C, Deckersbach T, Williams Z, Baker J T, Dougherty D D and Widge A S 2020 Case Report of Dual-Site Neurostimulation and Chronic Recording of Cortico-Striatal Circuitry in a Patient With Treatment Refractory Obsessive Compulsive Disorder *Front*. Hum. Neurosci. 14 569973

[36] Basu I, Robertson M M, Crocker B, Peled N, Farnes K, Vallejo-Lopez D I, Deng H, Thombs M, Martinez-Rubio C, Cheng J J, McDonald E, Dougherty D D, Eskandar E N, Widge A S, Paulk A C and Cash S S 2019 Consistent linear and non-linear responses to invasive electrical brain stimulation across individuals and primate species with implanted electrodes Brain Stimulat. 12 877–92

[37] Stypulkowski P H, Stanslaski S R, Jensen R M, Denison T J and Giftakis J E 2014 Brain Stimulation for Epilepsy – Local and Remote Modulation of Network Excitability Brain Stimulat. 7 350–8

[38] Cermak N, Wilson M A, Schiller J and Newman J P 2019 Stimjim: open source hardware for precise electrical stimulation 757716

[39] Siegle J H, López A C, Patel Y A, Abramov K, Ohayon S and Voigts J 2017 Open Ephys: an open-source, plugin-based platform for multichannel electrophysiology *J*. Neural Eng. 14 045003

[40] Arena A, Comolatti R, Thon S, Casali A G and Storm J F 2021 General Anesthesia Disrupts Complex Cortical Dynamics in Response to Intracranial Electrical Stimulation in Rats eNeuro 8

[41] Claar L D, Rembado I, Kuyat J R, Russo S, Marks L C, Olsen S R and Koch C 2023 Cortico-thalamo-cortical interactions modulate electrically evoked EEG responses in mice ed M B Eisen eLife 12 RP84630

[42] Panskus R, Holzapfel L, Serdijn W A and Giagka V 2023 On the Stimulation Artifact Reduction during Electrophysiological Recording of Compound Nerve Action Potentials * 2023 45th Annual International Conference of the IEEE Engineering in Medicine & Biology Society (EMBC) 2023 45th Annual International Conference of the IEEE Engineering in Medicine & Biology Society (EMBC) pp 1–5

[43] Kundu B, Davis T S, Philip B, Smith E H, Arain A, Peters A, Newman B, Butson C R and Rolston J D 2020 A systematic exploration of parameters affecting evoked intracranial potentials in patients with epilepsy Brain Stimulat. 13 1232–44

[44] Yamao Y, Matsumoto R, Kikuchi T, Yoshida K, Kunieda T and Miyamoto S 2021 Intraoperative Brain Mapping by Cortico-Cortical Evoked Potential *Front*. Hum. Neurosci. 15 635453

[45] Korhonen R J, Hernandez-Pavon J C, Metsomaa J, Mäki H, Ilmoniemi R J and Sarvas J 2011 Removal of large muscle artifacts from transcranial magnetic stimulation-evoked EEG by independent component analysis *Med. Biol*. Eng. Comput. 49 397–407

[46] Rogasch N C, Thomson R H, Farzan F, Fitzgibbon B M, Bailey N W, Hernandez-Pavon J C, Daskalakis Z J and Fitzgerald P B 2014 Removing artefacts from TMS-EEG recordings using independent component analysis: Importance for assessing prefrontal and motor cortex network properties NeuroImage 101 425–39

[47] Hyvarinen A 1999 Fast and robust fixed-point algorithms for independent component analysis *IEEE Trans*. Neural Netw. 10 626–34

[48] OPEN SCIENCE COLLABORATION 2015 Estimating the reproducibility of psychological science Science 349 aac4716

[49] Munafò M R, Nosek B A, Bishop D V M, Button K S, Chambers C D, Percie du Sert N, Simonsohn U, Wagenmakers E-J, Ware J J and Ioannidis J P A 2017 A manifesto for reproducible science Nat. Hum. Behav. 1 0021

[50] Landis S C, Amara S G, Asadullah K, Austin C P, Blumenstein R, Bradley E W, Crystal R G, Darnell R B, Ferrante R J, Fillit H, Finkelstein R, Fisher M, Gendelman H E, Golub R M, Goudreau J L, Gross R A, Gubitz A K, Hesterlee S E, Howells D W, Huguenard J, Kelner K, Koroshetz W, Krainc D, Lazic S E, Levine M S, Macleod M R, McCall J M, Moxley III R T, Narasimhan K, Noble L J, Perrin S, Porter J D, Steward O, Unger E, Utz U and Silberberg S D 2012 A call for transparent reporting to optimize the predictive value of preclinical research Nature 490 187–91

[51] Begley C G and Ioannidis J P A 2015 Reproducibility in Science *Circ*. Res. 116 116–26

[52] Fries P 2015 Rhythms For Cognition: Communication Through Coherence Neuron 88 220–35

[53] Cohen M X 2014 Analyzing Neural Time Series Data: Theory and Practice (Cambridge, MA: MIT Press)

[54] Muller L, Chavane F, Reynolds J and Sejnowski T J 2018 Cortical travelling waves: mechanisms and computational principles *Nat*. Rev. Neurosci. 19 255–68

[55] Morrow J K, Cohen M X and Gothard K M 2020 Mesoscopic-scale functional networks in the primate amygdala ed D Lee, K M Wassum and D Lee eLife 9 e57341

[56] Mitzdorf U 1985 Current source-density method and application in cat cerebral cortex: investigation of evoked potentials and EEG phenomena *Physiol*. Rev. 65 37–100

[57] Lindén H, Tetzlaff T, Potjans T C, Pettersen K H, Grün S, Diesmann M and Einevoll G T 2011 Modeling the Spatial Reach of the LFP Neuron 72 859–72

[58] Weerasinghe G, Duchet B, Bick C and Bogacz R 2021 Optimal closed-loop deep brain stimulation using multiple independently controlled contacts *PLoS Comput*. Biol. 17 e1009281

[59] Zelmann R, Paulk A C, Basu I, Sarma A, Yousefi A, Crocker B, Eskandar E, Williams Z, Cosgrove G R, Weisholtz D S, Dougherty D D, Truccolo W, Widge A S and Cash S S 2020 CLoSES: A platform for closed-loop intracranial stimulation in humans NeuroImage 223 117314

[60] Dastin-van Rijn E M, König S D, Carlson D, Goel V, Grande A, Nixdorf D R, Benish S, Widge A S, Nahas Z, Park M C, Netoff T I, Herman A B and Darrow D P 2022 Personalizing Dual-Target Cortical Stimulation with Bayesian Parameter Optimization Successfully Treats Central Post-Stroke Pain: A Case Report Brain Sci. 12 25

[61] Dastin-van Rijn E M and Widge A S 2025 Failure modes and mitigations for Bayesian optimization of neuromodulation parameters J. Neural Eng. 22 036038

[62] Nagrale S S, Yousefi A, Netoff T I and Widge A S 2023 In silico development and validation of Bayesian methods for optimizing deep brain stimulation to enhance cognitive control *J*. Neural Eng. 20 036015

